# Oncogenic Ras deregulates cell-substrate interactions during mitotic rounding and respreading to alter cell division orientation

**DOI:** 10.1101/2023.01.12.523730

**Authors:** Sushila Ganguli, Tom Wyatt, Tim Meyer, Buzz Baum, Helen K. Matthews

## Abstract

Oncogenic Ras has been shown to change the way cancer cells divide by increasing the forces generated during mitotic rounding. In this way, Ras^V12^ enables cancer cells to divide across a wider range of mechanical environments than normal cells. Here, we identify a further role for oncogenic Ras-ERK signalling in division by showing that Ras^V12^ expression alters the shape, division orientation and respreading dynamics of cells as they exit mitosis, in a manner that depends on MEK and ERK. Many of these effects appear to result from the impact of Ras^V12^ signalling on actomyosin contractility, since Ras^V12^ induces the severing of retraction fibres that normally guide spindle positioning and provide a memory of the interphase cell shape. In support of this idea, the Ras^V12^ phenotype is reversed by inhibition of actomyosin contractility, and can be mimicked by the loss of cell-substrate adhesion during mitosis. Thus, the induction of oncogenic Ras-ERK signalling leads to rapid changes in division orientation that, along with the effects of Ras^V12^ on cell growth and cell cycle progression, are likely to disrupt epithelial tissue organisation and contribute to cancer dissemination.

## Introduction

The Ras family of genes were among the first identified human oncogenes (Shih and Weinberg, 1982) and are the most frequently mutated in cancer (Sanchez-Vega *et al*., 2018). Activation of Ras oncogenes results in the hyperactivation of proliferative signalling pathways including the ERK and PI3K pathways to promote growth, survival and cell cycle entry (Krygowska and Castellano, 2018; Lavoie, Gagnon and Therrien, 2020). In addition, Ras-ERK signalling also affects cell shape and the organisation of the actin cytoskeleton (Helfman and Pawlak, 2005; Soriano *et al*., 2021). Ras-induced tranformation is therefore associated with changes in cell morphology and behaviours thought to aid invasion and metastasis (Pollock *et al*., 2005; Choi and Helfman, 2014; Logue *et al*., 2015; Mendoza *et al*., 2015). In recent work, we showed that oncogenic Ras-ERK signalling also impacts cell shape in mitosis (Matthews *et al*., 2020). Entry into mitosis is accompanied by cell rounding, driven by reorganisation of the actin cytoskeleton, loss of substrate adhesion and osmotic swelling (Taubenberger, Baum and Matthews, 2020). Mitotic cell rounding plays an important role in creating space for mitotic spindle formation and facilitates the subsequent alignment of the metaphase chromosomes Lancaster *et al*., 2013; Cadart *et al*., 2014). We previously showed that activation of oncogenic Ras-ERK signalling increased cell rounding in early mitosis. These changes in cell geometry were accompanied by alterations in cell mechanics that limit the DNA segregation errors observed in confined cell division (Matthews *et al*., 2020).

After rounding up at mitotic entry, cells then undergo a series of cell shape transitions as they divide. Cells elongate, and constriction of the cytokinetic ring in the centre of the cell divides them into two (Green, Paluch and Oegema, 2012). At the same time, the processes of de-adhesion and actomyosin remodelling that occurred during mitotic rounding are reversed as cells respread onto the substrate and re-establish their interphase shape. Following division, daughter cells respread along retraction fibres (Dix *et al*., 2018), thin acto-myosin rich fibres that anchor rounded mitotic cells to the substrate in mitosis (Cramer and Mitchison, 1993; Chen, Aretz and Kassler, 2022). Classical focal adhesion complexes are disassembled in early mitosis (Dix *et al*., 2018; Lock *et al*., 2018; Chen, Aretz and Kassler, 2022) but retraction fibres form at the location of previous focal adhesions and remain attached to the substrate by integrin-dependent adhesions during cell rounding, thus preserving a memory of the interphase cell shape. This enables daughter cells to re-occupy the interphase footprint of their mother during post-mitotic respreading (Mali, Wirtz and Searson, 2010; Dix *et al*., 2018).

Retraction fibres and substrate adhesion also affect positioning of the mitotic spindle which determines division orientation. In tissues, regulation of division orientation and post-mitotic cell shape is crucial for the appropriate positioning of daughter cells and therefore epithelial tissue architecture (Finegan and Bergstralh, 2019). The mitotic spindle is positioned with respect to cell shape and mechanical forces acting on the cell, including extracellular adhesive contacts with the substrate (Lechler and Mapelli, 2021). In single cells in adherent tissue culture conditions, retraction fibres play an important role in orienting the spindle (Théry *et al*., 2005; Toyoshima and Nishida, 2007; Fink *et al*., 2011) where they have been shown to exert force on the mitotic cell body to induce spindle rotation (Fink *et al*., 2011). In addition, integrin-mediated adhesion plays an important role in orienting the spindle in the plane of the substrate (Théry *et al*., 2005; Toyoshima and Nishida, 2007; Fink *et al*., 2011) to ensure both daughter cells remain attached to the substrate after division (Toyoshima and Nishida, 2007; Lechler and Mapelli, 2021).

As we had previously found that activation of Ras oncogenes can alter the dynamics of mitotic rounding, we wanted to investigate whether differences in mitotic exit were induced by Ras expression. We found that the expression of HRas that possesses the activating G12V mutation rapidly alters the division orientation, shape and respreading dynamics during mitotic exit. These effects result from the impact of Ras^V12^ on actomyosin contractility, since Ras^V12^ induces the severing of retraction fibres that normally guide spindle positioning and provide a memory of the interphase cell shape during accelerated mitotic rounding. In support of this idea, the Ras^V12^ phenotype is reversed by inhibition of actomyosin contractility, and can be mimicked by the loss of cell-substrate adhesion during mitosis. Together these findings show how activation of a single oncogene can directly alter the spatial control of cell division as a relatively early event in oncogenesis.

## Results

### Ras-ERK signalling affects the dynamics of post-mitotic respreading

To investigate the impact of short-term oncogenic Ras signalling we used the previously validated, tamoxifen-inducible oestrogen receptor (ER) HRas^V12^ fusion system (Molina-Arcas *et al*., 2013; Matthews *et al*., 2020) in the human, non-transformed epithelial cell line MCF10A (Soule *et al*., 1990). In this system, addition of 4-OH-tamoxifen (4-OHT) leads to rapid activation of HRasG12V (hereafter called Ras^V12^) and activation of ERK and Pi3K signalling over a period of 15 minutes to 24 hours (Molina-Arcas *et al*., 2013; Matthews *et al*., 2020). We previously found that five hours of Ras^V12^ activation induced changes in mitotic cell geometry by enhancing cell rounding in early mitosis (Matthews *et al*., 2020). We therefore wanted to follow these cells to see how they went on to divide. Bright-field time-lapse microscopy was used to follow unlabelled, asynchronous cells as they progressed beyond metaphase to divide and respread (Fig 1a). We measured metaphase and the combined daughter cell areas to assess respreading dynamics in the presence or absence of Ras^V12^ induction (Fig 1b). We found Ras^V12^ activation accelerated the rate of post-mitotic respreading (Fig 1b) and resulted in a significant increase in respread cell area within 10 minutes of the onset of anaphase (Figure 1c).

**Figure 1.**
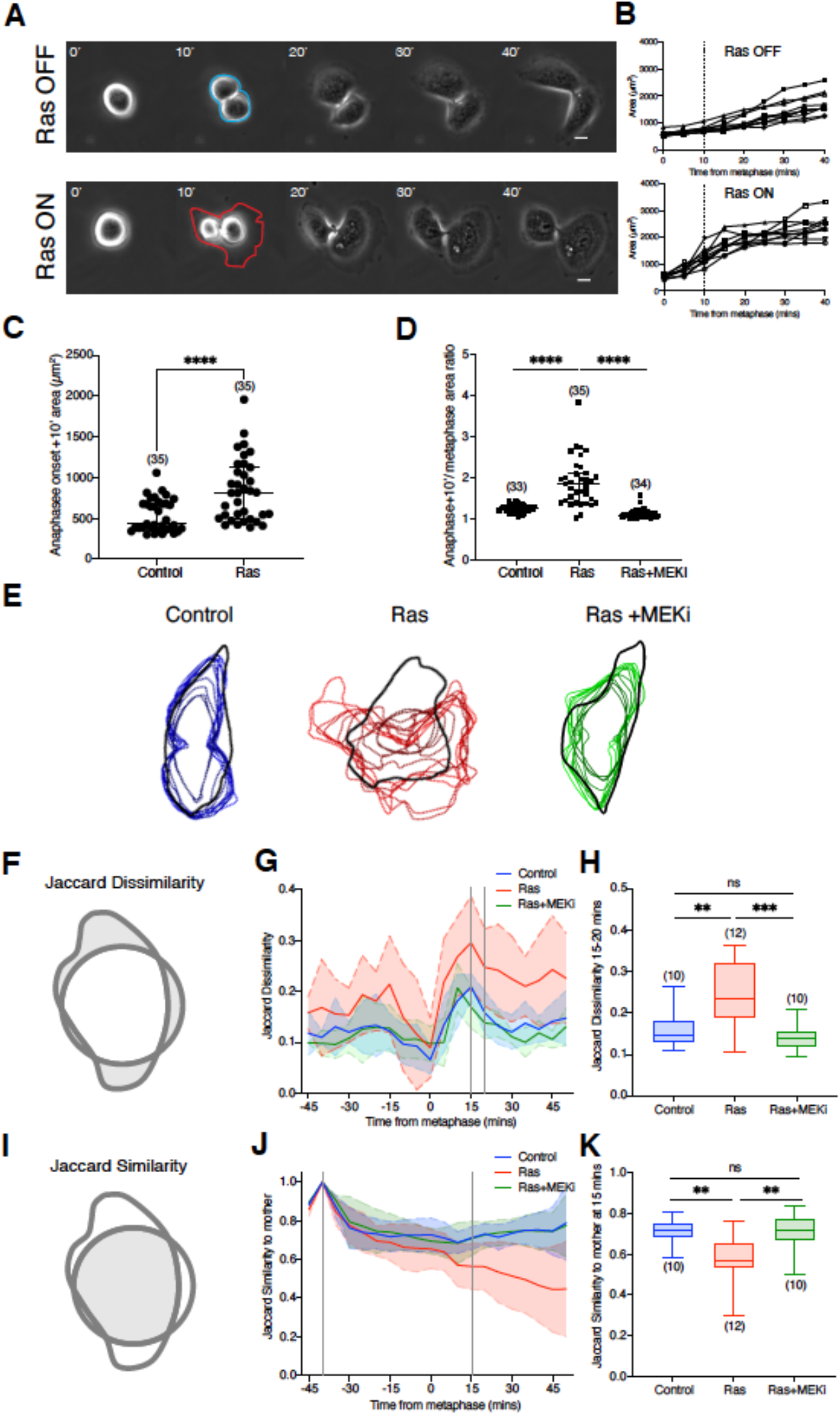
Ras-ERK signalling affects the dynamics of post-mitotic respreading. (A) Representative phase contrast images of ER-Ras^V12^ cells exiting division following ethanol or 4-OHT treatment. Time in minutes is aligned so that t = 0 is the frame 5 minutes before the first evidence of anaphase elongation or visible chromosome separation. Metaphase and combined daughter cell areas were manually segmented from images as illustrated; ethanol (blue) and 4-OHT (red). Scale bars represent 10μm. (B) Quantification of cell area for 10 ER-Ras^V12^ cells exiting mitosis following ethanol or 4-OHT treatment, as described in (A). Measurements were taken from phase-contrast time-lapse microscopy images of cells every 5 minutes following 5–15 h of treatment. Dotted lines indicate 10 minutes after the onset of anaphase from which stastistical analysis is calculated in (C). (C) Plot of cell area 10 minutes following the onset of anaphase of ER-Ras^V12^ cells treated with ethanol or 4-OHT. N= 3 experiments. (D) Plot of the ratio of cell area (10 minutes following anaphase onset/ metaphase) for ER-Ras^V12^ cells following ethanol, 4-OHT or 4-OHT + 10 μM selumetinib (MEKi) treatment. N= 3 experiments. (E) Cell outlines as cells respread post-mitosis with respect to the mother cell shape. MCF10A-LifeAct-GFP-ER-Ras^V12^ cells were treated with ethanol, 4-OHT or 4-OHT plus 10μM selumetinib for 5-15 hours prior to time-lapse bright-field and fluorescence imaging at 5 minute intervals. Representative dividing cells were manually segmented at interphase (black), defined using the bright-field channel as 15 minutes before nuclear envelope breakdown (NEB). Post-mitotic respreading cell outlines were segmented at 5-minute intervals for 10 consecutive frames from the final frame of metaphase following ethanol (blue), 4-OHT (red) or 4-OHT plus MEKi (green). Image overlays were produced using CellProfiler software as described in Methods. (F) Diagram showing the definition of the Jaccard dissimilarity measurement. Grey indicates the non-overlapping area of two cell shapes. (G) Graph to show the Jaccard dissimilarity measurement (normalised for area and centroid displacement) as cells progress through mitosis comparing cell shapes between consecutive time-points (5-minute intervals). Dotted lines indicate 15 and 20 minutes following anaphase onset, used for statistical analysis in (G). n= 10 cells in each condition. (H) Box-whisker plot to show the Jaccard dissimilarity between 15- and 20-minute time-points following anaphase onset for all three conditions. (I) Diagram to show the definition of the Jaccard similarity measurement. Grey indicates the overlapping area of two cell shapes. (J) Graph to show the Jaccard similarity (normalised for area and centroid displacement) of cells as they progress through mitosis at consecutive time-points compared with the mother cell. Dotted lines indicates 15 minutes before NEB and 15 minutes following the onset of anaphase, used for statistical analysis in (K). n= 10 cells in each condition. (K)Box-whisker plot to show the Jaccard similarity comparing timepoints 15 minutes following anaphase onset with the mother cell for all three conditions.

As the effects of Ras^V12^ on mitotic rounding are mediated through ERK signalling, we next tested whether the change in rate of post-mitotic respreading was sensitive to MEK inhibition. For this analysis, inducible ER-Ras^V12^ cells were treated with ethanol, 4-OHT or 4-OHT plus the previously validated MEK inhibitor selumetinib (Matthews *et al*., 2020). As MEK inhibition reduces the rate of mitotic rounding (Matthews *et al*., 2020), a comparison of absolute cell areas could artificially increase post-metaphase respreading due to the larger starting metaphase cell area. We therefore normalised the analysis by calculating the ratio of the cell area at metaphase with the area 10 minutes after anaphase-onset (post-anaphase cell area/ metaphase cell area). In this way, we observed a significant increase in the rate of respreading in early anaphase following Ras^V12^ expression, which was reversed with inhibition of ERK signalling (Fig 1d). Taken together with the cell shape changes at mitotic entry, these data show that Ras-ERK signalling increases the rate at which cells change their shape during both mitotic rounding and post-mitotic cell respreading.

While studying the dynamics of post-mitotic respreading, we observed that Ras^V12^ expression also altered the shape of cells as they respread. To examine this phenotype in more detail, we used LifeAct-GFP labelled cells to enhance tracking of the cell margin (S1a). In this experiment, ER-Ras^V12^-LifeAct-GFP cells were treated with ethanol, 4-OHT or 4-OHT plus selumetinib immediately prior to live bright-field and fluorescence imaging for 15 hours. We segmented cells at interphase and as they exited mitosis, and the GFP-labelled cell contours were overlaid (Fig 1e and S1b).

This indicated that while control cells and cells treated with 4-OHT+MEK inhibitor maintained a consistent shape as they exited mitosis, the shape of cells expressing oncogenic Ras^V12^ during post-mitotic respreading appeared variable - indicating that Ras-ERK signalling has a dramatic impact on the dynamics of cell shape changes that accompany mitotic exit. As a simple quantitative measure of shape similarity to analyse respreading dynamics, we used the Jaccard index (Jaccard, 1912). This measures the relative overlap of different shapes (Fig 1i). It can also be extended to define a Jaccard dissimilarity index as the area of the non-overlapping regions of the two shapes (Fig 1f). Changes in Jaccard dissimilarity can arise from changes in cell area or displacement as well as cell shape, and we found that both area change and displacement are increased by oncogenic Ras (Fig 1c and S1c, d). Therefore, in order to limit this measure to changes in cell shape, the contribution of area and displacement was removed by scaling and translating the images (see Methods). This analysis demonstrated that despite the contributions of area change and displacement, Ras^V12^ activation increases shape dynamics during post-mitotic respreading (Fig 1g) and significantly increases the rate at which cells change shape as they exit mitosis (Fig 1h).

In cell culture models of division, daughter cells tend to take up the shape of the mother cell footprint (Théry and Bornens, 2006). Previous work has shown that this depends on the integrin-ECM adhesions and retraction fibres that persist during mitosis (Cramer and Mitchison, 1993) acting as a physical memory of interphase shape and guiding daughter cell respreading (Dix *et al*., 2018; Lock *et al*., 2018). In line with these previous studies, we demonstrated that control and MEK inhibited cells adopt the same shape as the mother cell as they respread (Fig 1e). By contrast, cells expressing Ras^V12^ tended to escape the confines of the mother cell footprint. We quantified these effects using the Jaccard similarity index (Fig 1i), adapted to remove changes in area and displacement. As control and MEK inhibited cells started to exit mitosis, the Jaccard similarity increased reflecting their ability to take up the mother cell shape (Fig 1j). However, over the same period, we observed a significant decrease in the similarity of Ras^V12^ expressing cells with the mother cell footprint (Fig 1k). Taken together, these data show that short-term expression of oncogenic HRAS increases the dynamics of cell spreading in cells leaving mitosis and decreases the likelihood of daughter cells taking up their mother cell area.

### Ras-ERK signalling induces the asymmetric respreading of daughter cells

Through studying the dynamics of cell respreading during mitotic exit, we also observed that Ras^V12^ expression frequently causes differences in the rate of respreading between daughter cells from the same division. To quantify this change in symmetry, bright-field imaging was used to follow unlabelled, asynchronous ER-Ras^V12^ cells treated with ethanol or 4-OHT immediately prior to imaging as they exited mitosis and divided (Fig 2a). Daughter cell areas were manually segmented at 20 minutes following anaphase onset - a time when respreading has mostly been completed (Fig 1b and S1c). As a measure of the symmetry of respreading, the ratio of the two daughter cell areas (larger/smaller) was calculated and plotted against time (Fig 2b). While ethanol-treated control cells tended to divide symmetrically over 23 hours of imaging, maintaining a ratio of <2 in 97.5% of divisions (Fig 2b, left-hand graph), 37% of cells expressing Ras^V12^ underwent asymmetric divisions (ratio≥2) (Fig 2b, right-hand graph). Remarkably, this phenotype was observed as early as 3 hours after 4-OHT addition.

**Figure 2.**
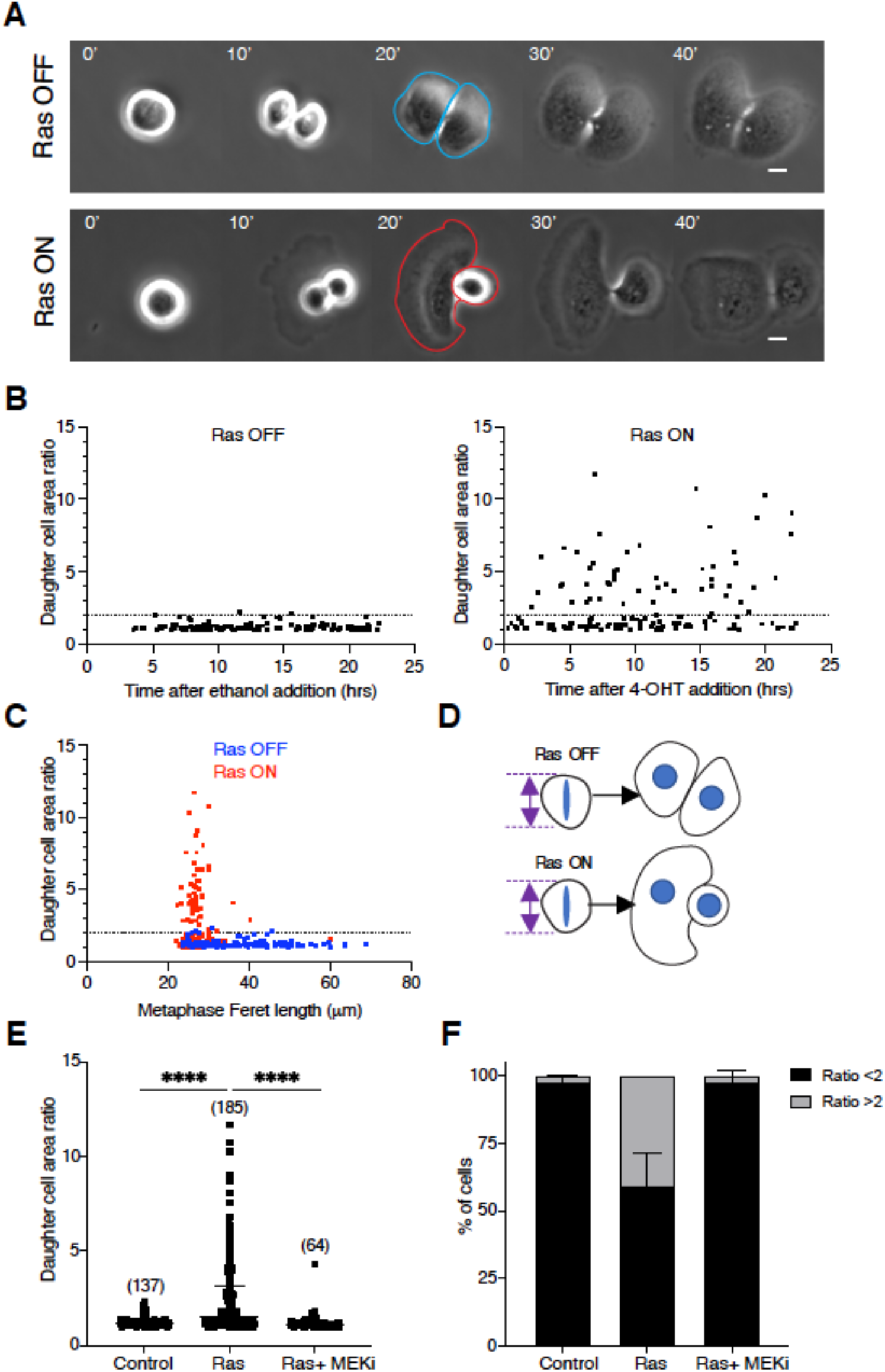
Ras-ERK signalling induces the asymmetric respreading of daughter cells. (A) Representative phase contrast images of ER-Ras^V12^ cells exiting division following ethanol or 4-OHT treatment. Cells were imaged following 5-15 h of treatment at 5 minute intervals. Time in minutes is aligned so that t = 0 is the frame 5 minutes before evidence of anaphase elongation or visible chromosome separation. Individual daughter cell areas are segmented from images as illustrated; ethanol (blue) and 4-OHT (red). Scale bars represent 10μm. (B) Plot of the daughter cell area ratio of individual ER-Ras^V12^ cells dividing against time after ethanol addition (left-hand graph) or 4-OHT addition (right-hand graph). Measurements of individual daughter cell areas were taken at 20 minutes following the onset of anaphase. The ratio was measured as larger/smaller. Dotted line indicates daughter cell area ratio of 2. N= 2 experiments. (C) Plot of the metaphase cell length (Feret) against daughter cell area ratio of individual ER-Ras^V12^ cells dividing following ethanol (blue) or 4-OHT (red) treatments. Dotted line indicates daughter cell area ratio of 2. N= 2 experiments. (D) Diagram depicting how control cells that are able to round up at metaphase to the same extent as Ras^V12^ activated cells continue to divide symmetrically. (E) Plot of daughter cell area ratio of ER-Ras^V12^ cells following ethanol, 4-OHT or 4-OHT plus selumetinib treatment. Measurements were taken from phase contrast imaging of ER-Ras^V12^ cells as described in (A). N= 3 experiments. (F) Graph showing the percentage of cells from the data in (E) that divide with a daughter cell area ratio of < or ≥ 2. Error bars show mean and SD. N= 3 experiments.

Since we previously demonstrated that Ras signalling affects mitotic rounding (Matthews *et al*., 2020), we then asked whether asymmetric respreading was due to the fact that Ras^V12^-activated cells were rounder in mitosis by plotting the corresponding metaphase cell length of each dividing cell against the resulting daughter cell area ratio (Fig 2c). This showed that asymmetric divisions were frequently seen in Ras^V12^ expressing cells, but rarely in control cells that were also able to round up. This suggests that while Ras^V12^ expression induces both accelerated mitotic rounding and asymmetric respreading, the change in division symmetry cannot be explained by the impact on mitotic cell shape alone (Fig 2d). Importantly, this effect of Ras^V12^ on division symmetry was abolished by treatment with the MEK inhibitor (Fig 2e, f).

### Activation of oncogenic Ras causes misalignment of the mitotic spindle

Asymmetric respreading could arise as a consequence of alterations in division orientation as the plane of cell division in animal cells is determined by the orientation of the mitotic spindle (Lechler and Mapelli, 2021). We tested this idea by constructing MCF10A ER-Ras^V12^ cell lines that stably express tubulin-GFP to visualise mitotic spindle dynamics during mitosis. Confocal live cell imaging was then used to follow tubulin-GFP in ER-Ras^V12^ cells treated with ethanol or 4-OHT as they divided and respread (Fig 3a). As observed in XZ cross-sections, the spindle and mid-body of Ras^V12^ activated cells were frequently aligned obliquely with respect to the substrate, so that only one of the two daughter cells was in contact with the substrate and able to respread rapidly upon exit from mitosis (Fig 3a, b).

**Figure 3.**
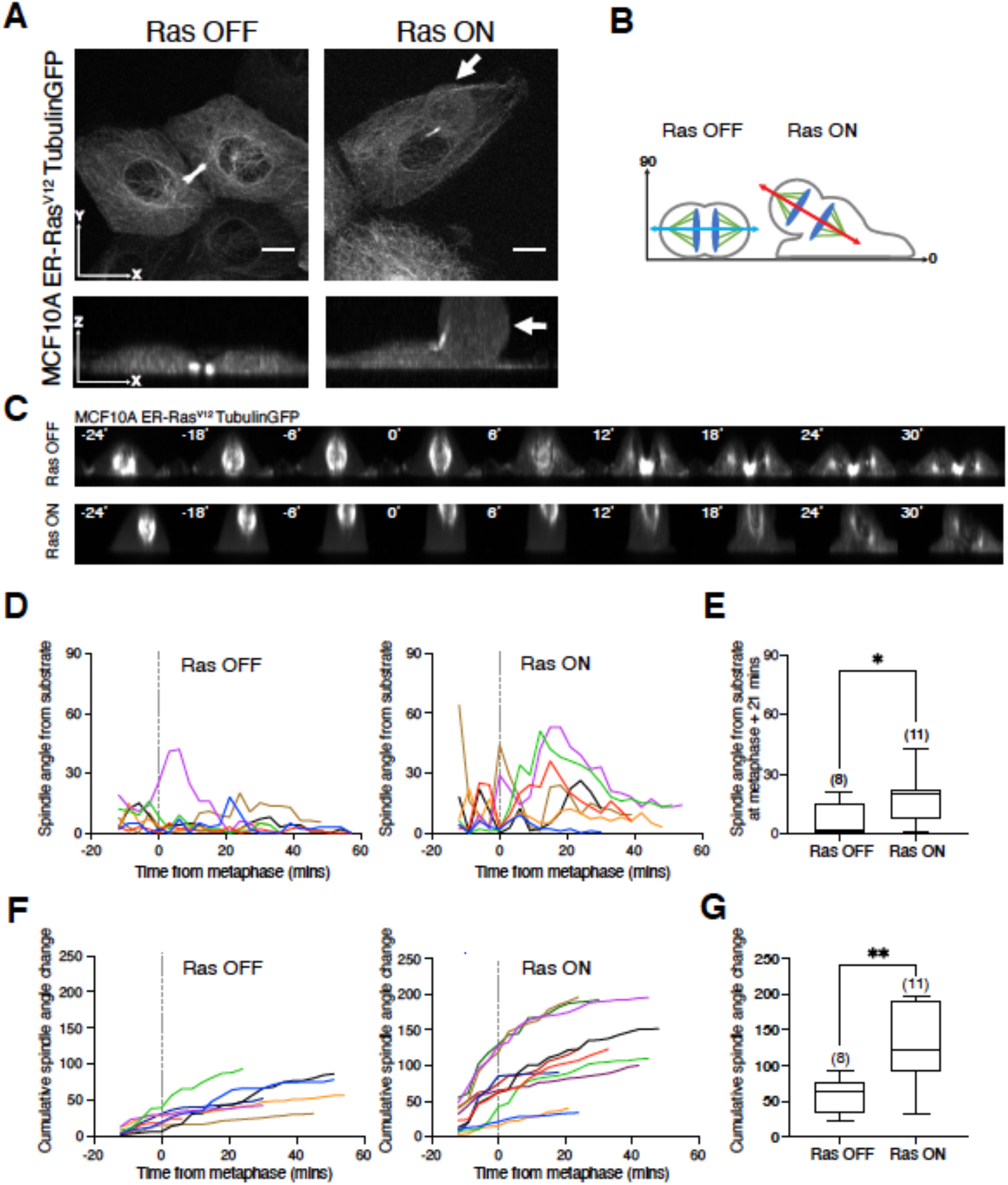
Activation of oncogenic Ras causes the misalignment of the mitotic spindle. (A) Representative images of MCF10A ER-Ras^V12^ cells labelled with LifeAct-tubulin following a 5 h treatment with ethanol or 4-OHT. Images are displayed as maximum projections in x-y and x-z. Arrows indicate an apically positioned daughter cell. Scale bars represent 10 μm. (B) Schematic to show spindle alignment relative to the substrate according to Ras^V12^ activation. (C) Representative time-lapse images of MCF10A ER-Ras^V12^ cells labelled with tubulin-GFP progressing through mitosis following a 5-hour treatment of ethanol or 4-OHT taken at 3 minute intervals. Images are presented as maximum projections in x-z. Time is aligned so t=0 represents the frame 3 minutes before anaphase spindle elongation. Manual segmentation of the axis between the two spindle poles from the time of the first appearance of the mitotic spindle enabled calculation of the angle of the spindle relative to the substrate (x axis = 0 degrees). (D) Quantification of the spindle angle relative to the substrate as described in (C) over time for 7 individual MCF10A ER-Ras^V12^-tubulinGFP cells following a 5-hour treatment of ethanol or 4-OHT. Dotted lines at t=0 represents the frame 3 minutes before anaphase spindle elongation. (E) Box-whisker plot to show the spindle angle at 21 minutes following the onset of anaphase for ER-Ras^V12^-tubulinGFP cells following a 5-hour treatment of ethanol or 4-OHT. (F) Graphs to show the cumulative spindle angle from substrate change as cells progress through mitosis at 3 minute intervals for individual cells as described in (C). (G) Box-whisker plot to show the cumulative spindle angle change for ER-Ras^V12^-tubulinGFP cells following a 5-hour treatment of ethanol or 4-OHT.

To further characterise the impact of Ras^V12^ activation on the positioning of the mitotic spindle in the plane of the substrate, live cell imaging was again used to follow tubulin-GFP labelled spindles in ER-Ras^V12^ cells treated with ethanol or 4-OHT as they progressed through mitosis. To reveal the angle of the spindle relative to the substrate, we generated cross-sectional images (Fig 3c). This showed that spindles in Ras^V12^ activated cells tended to have an unstable position and were frequently tilted at an angle of >30 degrees from the substrate (Fig 3d). While some control cells transiently exhibited titled spindles (>30 degrees), such errors tended to be corrected soon after mitotic exit, resulting in interphase daughter cells positioned next to one another on the substrate within 21 minutes of anaphase (Fig 3e). By contrast, cells expressing Ras^V12^ exhibited large defects in spindle orientation early in mitosis (>30 degrees), and took much longer to correct these defects during mitotic exit (Fig 3d, e). The combined effects of these processes led to a significant increase in the cumulative movement of spindles in Ras^V12^-activated cells relative to the control (Fig 3f, g).

### Ras activation induces breakages in retraction fibres

To determine the underlying mechanism of this Ras^V12^-induced change in spindle orientation, we examined cell substrate adhesions. Although adhesions are remodelled as cells pass into and out of mitosis (Ramkumar and Baum, 2016), mitotic cells maintain physical connections with the extra-cellular environment through retraction fibres (Cramer and Mitchison, 1993; Chen, Aretz and Kassler, 2022). These structures, formed during mitotic rounding, are tethered to the substrate at their tips by integrin-based structures that lack many of the components associated with interphase adhesions (Dix *et al*., 2018; Lock *et al*., 2018). Importantly, these mitotic adhesive structures have also been implicated in positioning of the mitotic spindle (Théry *et al*., 2005; Fink *et al*., 2011) and have been shown to guide post-mitotic respreading (Dix *et al*., 2018; Lock *et al*., 2018). When we imaged retraction fibres in fixed metaphase cells (Fig 4a and S2), we frequently observed retraction fibres that appeared to be severed in Ras^V12^-activated cells (Fig 4a). The proportion of cells with severed retraction fibres was significantly higher in Ras^V12^ activated cells compared to controls or cells treated with a MEK inhibitor (Fig 4b, c). This suggests the hypothesis that the effects of Ras^V12^ on mitotic exit could be mediated by the breakage of retraction fibres severing the cell’s connection with the substrate. To test this idea, we carried out an experiment to determine whether the loss of substrate attachment during mitosis might reproduce the effects of Ras^V12^ on spindle orientation and daughter cell respreading. In this experiment, mitotic cells treated with ethanol or tamoxifen for 15 hours were dislodged by mechanical force into suspension before being replated and immediately imaged exiting mitosis (Fig 4d and S4a).

**Figure 4.**
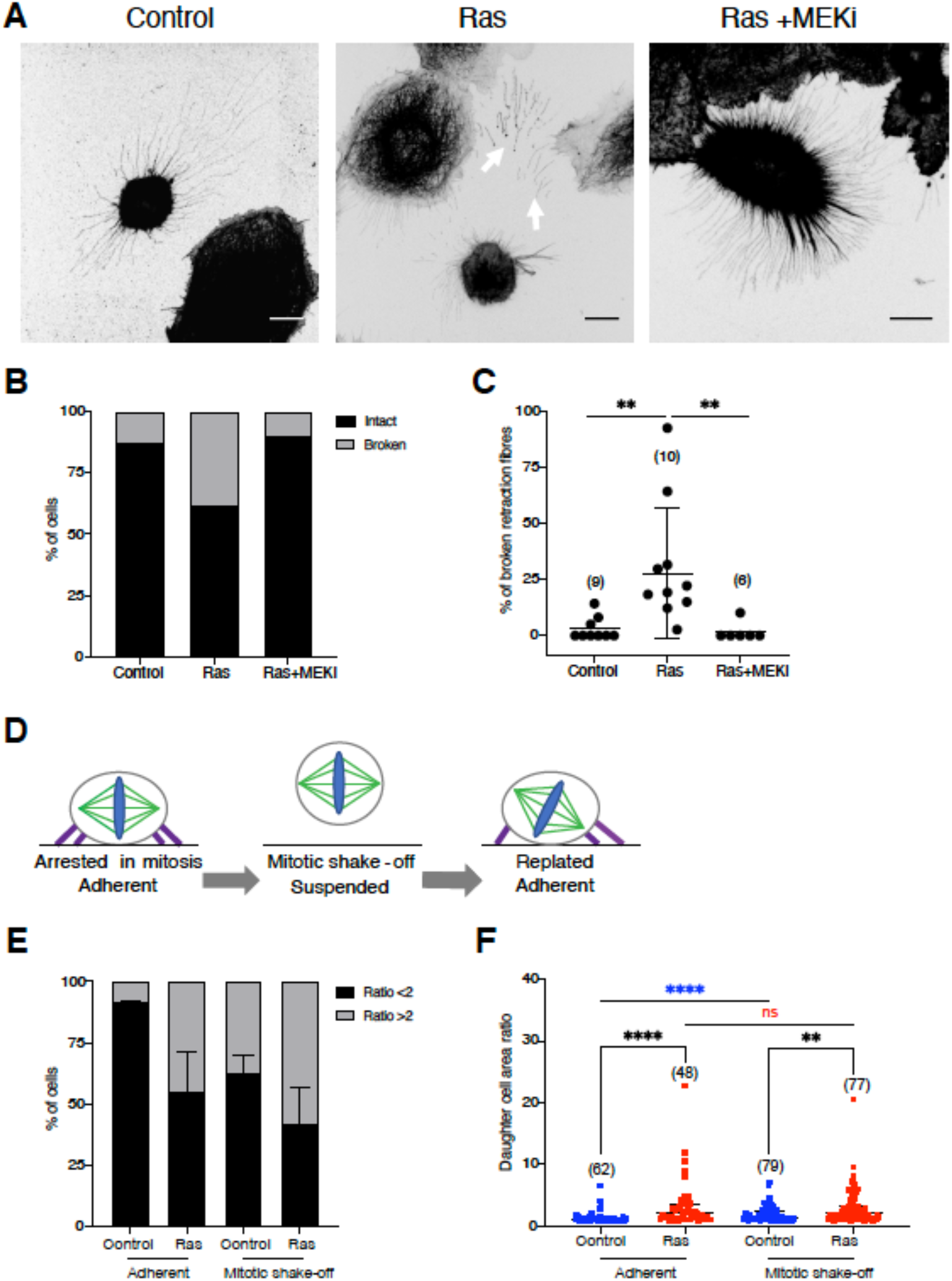
Activation of oncogenic Ras induces breakages in retraction fibres. (A) Immunofluorescence images of MCF10A-ER-Ras^V12^ cells in metaphase following a 5-hour treatment of ethanol, 4-OHT or 4-OHT+ 10 μM selumetinib (MEK inhibitor). Cells are stained for monomeric α-tubulin. Images are displayed as maxiumum projections. Arrows indicate breakages in retraction fibres. Scale bars represent 10μm. (B) Graph showing the percentage of cells as described in (A) with evidence of any broken retraction fibres. n = control (15), Ras (19), Ras+MEKi (10). (C) Quantification of the percentage of broken retraction fibres in single cells from the conditions described in (A). (D) Schematic to illustrate the mitotic shake-off protocol. ER-Ras^V12^ cells were treated with 10μmol STLC and ethanol or tamoxifen for 15 hours prior to ‘mitotic shake-off’. Mitotic cells are then dislodged by mechanical force into suspension. Upon removal of STLC, cells are then replated onto fibronectin -coated glass dishes in ethanol or tamoxifen-containing media and immediately imaged using bright-field time-lapse microscopy at 5 minute intervals as they exit mitosis. (E) Graph showing the percentage of cells that divide with a daughter cell area ratio of < or ≥ 2 according to whether they are subjected to mitotic shake-off or not. Adherent cells are plated on fibronectin-coated glass dishes for 24 hours, and treated with ethanol or 4-OHT immediately prior to imaging. Mitotic shake-off cells were treated as described in (D). Measurements of daughter cell areas at 20 minutes following the onset of anaphase were taken for individual cells and the percentage of cells with a daughter cell area ratio of < or ≥ 2 was calculated. Error bars show mean and SD. N=3 experiments. (F) Individual daughter cell area ratio measurements from the experiment described in (E).

To assess the effect of disrupting mitotic substrate adhesion on mitotic exit, we then measured the daughter cell area ratio at 20 minutes following anaphase onset. Control cells replated from suspension displayed a marked increase in division asymmetry (Fig 4e), which was accompanied by a significant increase in daughter cell asymmetry relative to cells that were not subject to mitotic shake-off (Fig 4f) - as expected if these cells are less able to determine the substrate plane. Many of these defects in spindle orientation were corrected as control cells exited mitosis and respread (Fig 4f). By contrast, Ras^V12^-activated cells exhibited similar levels of spindle mis-orientation and division asymmetry whether or not they were removed from the substrate during mitosis (Fig 4e, f). Furthermore, cells expressing oncogenic Ras were unable to correct any spindle defects as they respread (Fig 4f). Taken together, these data demonstrate that while the maintenance of mitotic cell-substrate interactions is crucial for correct spindle orientation and symmetrical respreading in normal cells grown on a substrate, the ability of cells to read the substrate in this way is disrupted by the expression of oncogenic Ras.

### The effects of oncogenic Ras on mitotic exit depend on acto-myosin contractility

Since oncogenic Ras induces actomyosin contractility to drive mitotic rounding (Matthews *et al*., 2020), we tested whether retraction fibres in Ras^V12^-activated cells are broken as a result of actomyosin contractility. To do this, we imaged Ras^V12^-expressing cells dividing in the presence or absence of an inhibitor of Rho kinase (ROCK), as ROCK and actomyosin contractility have a previously well described role in generating force during mitotic rounding (Maddox and Burridge, 2003; Matthews *et al*., 2012) (Fig S4b). Treatment with ROCK inhibitor prevented the severing of retraction fibres (Fig 5a, b and S3a) and increased the total number of retraction fibres (Fig S3b). ROCK inhibitor treatment restored normal division orientation and post-mitotic respreading in Ras^V12^-activated cells (Fig 5c, d). Finally, we also assessed whether inhibiting contractility in Ras^V12^-activated cells could restore the respreading of daughter cells into the mother cell footprint, as measured by the Jaccard similarity index (Fig 5e). Again, the effects of oncogenic Ras were reversed upon treatment with the ROCK inhibitor (Fig 5f). These data demonstrate that acto-myosin contractility is crucial for the effects of Ras^V12^ activation on cell division orientation and post-mitotic respreading.

**Figure 5.**
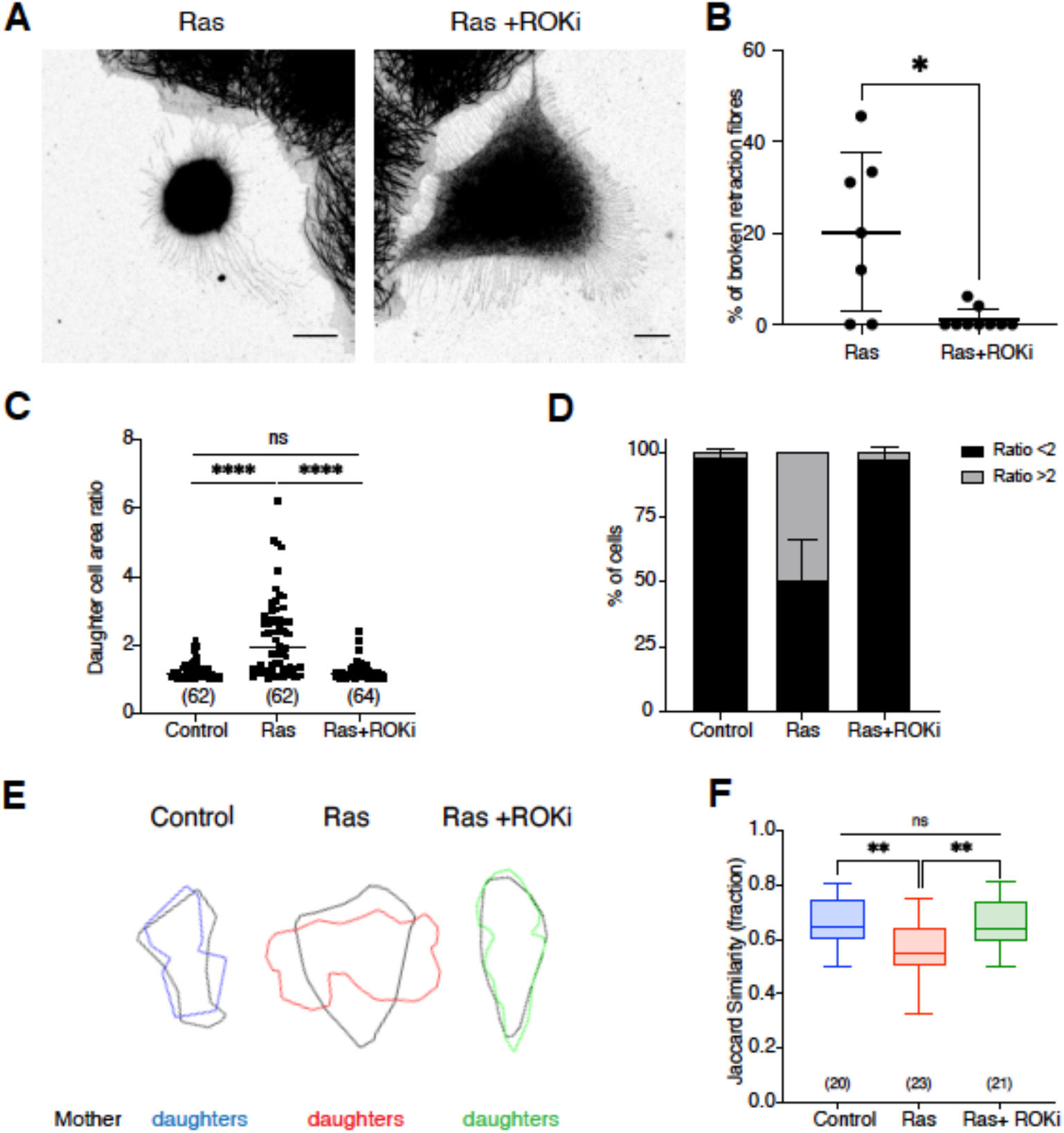
The effects of oncogenic Ras on mitotic exit depend on actomyosin contractility. (A) Immunofluorescence images of MCF10A-ER-Ras^V12^ cells in metaphase following a 5-hour treatment of 4-OHT or 4-OHT+ 25μM Y27632 (ROCK inhibitor). Cells are stained for monomeric α-tubulin. Images are displayed as maximum projections. Scale bars represent 10μm. (B) Quantification of the percentage of broken retraction fibres in single cells from the conditions described in (A). n= Ras (7), Ras+ROKi (9). (C) Quantification of daughter cell area ratio for ER-Ras^V12^ cells following ethanol, 4-OHT or 4-OHT plus ROCK inhibitor treatment. Measurements taken from bright-field time-lapse microscopy images as previously described. N= 3 experiments. (D) Graph showing the percentage of cells from the data in (C) that divide with a daughter cell area ratio of < or ≥ 2. Error bars show mean and SD. N= 3 experiments. (E) Overlay of mother and daughter cell outlines. Representative cells from the experiment described in (C) were manually segmented at NEB minus 15 minutes (mother) and 15 minutes following the onset of anaphase (daughters). (F) The Jaccard similarity index (shape contribution only) quantifying the overlap between mother and daughter cell shapes as defined in (E).

## Discussion

The regulation of cell division is crucial for the maintainence of tissue architecture and organisation (Lechler and Mapelli, 2021). In this study, we find that expression of the oncogene HRas^G12V^ alters substrate adhesion during mitosis, resulting in misregulation of cell division orientation and post-mitotic respreading. We previously found that Ras^V12^ activation increases acto-myosin contractility and allows cells to generate more force during cell rounding in early mitosis (Matthews *et al*., 2020). The new data presented here support a model by which this increased rounding force in early mitosis can act to sever retraction fibres that anchor mitotic cells to the substrate. This loss of substrate cues during division has two major effects: firstly, it leads to spindle misorientation relative to the substrate and, secondly, it prevents daughter cells reoccupying the footprint of their mother following division. Remarkably these effects of deregulated Ras signalling occur within a very short space of time following Ras^V12^ expression, and can be rapidly reversed by MEK inhibitors. While these changes could be due to Ras-ERK-mediated changes in transcription, the speed of these changes suggest the possibility of this being a more direct effect of signalling, as has been suggested by other studies implicating a direct role for ERK on acto-myosin contractility (Helfman and Pawlak, 2005; Mendoza *et al*., 2011, 2015). It remains to be determined which are key here.

This work is focused on the effects of Ras^V12^ activation in single cells where interactions with the extracellular matrix (ECM) are unidirectional and fairly well defined. In such systems, integrin and actin-rich retraction fibres have been shown to maintain substrate attachment following mitotic rounding to guide spindle alignment (Théry *et al*., 2005; Fink *et al*., 2011) as well as post-mitotic respreading (Cramer and Mitchison, 1993; Dix *et al*., 2018). Precisely how cell-substrate adhesions regulate division orientation is not completely clear. The forces required for spindle orientation are thought to depend on interactions between astral microtubules and the cell cortex (Lechler and Mapelli, 2021), which is patterned by polarity factors that integrate multiple cues from substrate adhesion, cell shape and chromatin (Kiyomitsu and Cheeseman, 2012; Dimitracopoulos *et al*., 2020). This is likely to apply to our system, and our data suggests that by compromising cell-substrate adhesion, Ras^V12^ activation interrupts the transmission of contractile forces along retraction fibres. This has the effect of impairing the normal process of mitotic spindle orientation and results in defects in daughter cell placement following mitotic exit.

It is not clear how ECM attachment is modulated during cell division in 3D tissues. While pseudo-stratified epithelia have been shown to maintain attachment to the basal lamina during mitotic rounding via thin basal processes which direct cleavage plane orientation (Kosodo *et al*., 2008; Nakajima *et al*., 2013), the extent to which these retraction fibres contribute to division orientation in tissues where additional cell-to-cell adhesions and polarity complexes are also at play remains to be determined (Bosveld *et al*., 2016). Changes to division orientation in this context have important implications, as they facilitate stress relaxation, maintain cell packing (Wyatt *et al*., 2015; Lisica *et al*., 2022) and epithelial architecture (Lechler and Mapelli, 2021). While errors in division orientation are tolerated in some tissues (Bergstralh, Lovegrove and St. Johnston, 2015), departure from planar divisions can disrupt epithelial organisation (Zheng *et al*., 2010; Bergstralh, Dawney and St Johnston, 2017), leading to the idea that spindle misorientation might contribute to oncogenesis (Macara and Seldin, 2017). Several studies support this hypothesis by showing how disruption of planar spindle alignment can lead to cell delamination (Nakajima *et al*., 2013), the formation of tumour-like masses (Quyn *et al*., 2010; Nakajima *et al*., 2013), as well as invasive cancer (Caussinus and Gonzalez, 2005). In addition, there is an association between spindle misorientation phenotypes and oncogenic signalling (Toyoshima *et al*., 2007; Fleming *et al*., 2009; Quyn *et al*., 2010; Vitiello *et al*., 2014). As the activation of Ras oncogenes have been shown to disrupt tissue structure inducing 2D-3D transitions in tissue culture (Nyga *et al*., 2021) and form tumour-like structures *in vivo* (Moruzzi *et al*., 2021; Nyga *et al*., 2022), it will be important to assess how far altered cell division contributes to loss of tissue structure in 3D models, and whether these early consequences of oncogene signalling might contribute to tumour formation and spread.

Our findings reveal a mechanism by which Ras mutations affect cell division orientation and post mitotic shape changes, which have the potential to impact tissue organisation during the earliest stages of tumour formation. This study has focused on HRAS^G12V^, a driver mutation in several cancers including bladder and thyroid cancer and squamous cell carcinoma (Hobbs, Der and Rossman, 2016), where early HRAS^G12V^ mutations drive abnormal tissue growth and folding (Fiore *et al*., 2020). In our previous study, we found similar mitotic rounding phenotypes following oncogenic KRAS and HRAS expression (Matthews *et al*., 2020). It will therefore be important future work to test whether our model may be more widely applicable to cancers driven by different Ras isoforms. Taken together, we show that Ras oncogenes act via ERK to alter cell contractility in a way that impacts the mechanics and outcome of cell division. The consequences of these are likely be profound when combined with the influence of Ras on EMT, cell growth and cell cycle progression. How a single pathway is able to influence such a breadth of cell biology is unclear, but these multiple effects of Ras-ERK signalling on growth and division may explain why Ras oncogenes are such potent drivers of cancer development and progression.

## Methods

### Cell lines and culture

MCF10A-ER-Ras^V12^ cell lines (gifts from J. Downward, Francis Crick Institute, London, UK) (Molina-Arcas *et al*., 2013) were cultured in phenol-free DMEM F-12 Glutamax with 5% charcoal-stripped horse serum (Invitrogen), 20ng/ml EGF (Peprotech), 0.5mg/ml Hydrocortisone (Sigma), 100ng/ml Cholera toxin (Sigma), 10*μ*g/ml Insulin (Sigma), 1% Penstrep (Gibco) at 37°C with 5% CO_2._ LifeAct-GFP labelled lines were produced by infection with puromycin resistant lentivirus (rLV-Ubi-LifeAct-GFP2, Ibidi 60141). Tubulin-GFP labelled lines were produced through infection with puromycin resistant retrovirus. GFP positive cells were sorted using flow cytometry at the FACS facility at UCL Great Ormond Street Institute of Child Health to produce a polyclonal stable pool.

### Drug treatments

Ras^V12^ was activated in inducible lines by addition of 100nM 4-OH-tamoxifen (Sigma). The following small molecule inhibitors were used in this study: MEK inhibitor: 10*μ*M Selumetinib (Selleckchem), ROCK inhibitor: 25*μ*M Y27632 (Sigma). Where indicated, control treatments were performed with equivalent amounts of ethanol. Details of treatment times are described in the figure legends.

To synchronise cells in metaphase for mitotic shake-off, cells were incubated with 10μmol STLC (Sigma) for 15 hours. STLC was then washed out, using 3 washes, before replacing with fresh media as described in the figure legends.

### Live cell imaging

For live cell microscopy, cells were plated in fibronectin-coated, glass-bottomed plates (Mattek). Cells were plated 24 hours before imaging with the exception of the mitotic shake-off experiments where they were replated immediately before imaging. Wide field, brightfield time-lapse imaging at 37°C was carried out on a Nikon Ti inverted microscope at 5-minute intervals using a 20x (Plan Fluor ELWD Ph1 NA 0.45, WD 7.4) or 40x (Plan Fluor ELWD Ph2 NA 0.6, WD 3.7) objective. Live confocal imaging at 37°C was carried out on the 3i spinning disc confocal microscope using the 63x oil objective (Plan Apochromat NA 1.4, WD 0.19), with 1μm z-steps at 3-minute intervals.

### Cell fixation and immunostaining

For immunofluorescence imaging, cells were plated 24 hours prior to instituting the experimental conditions and subsequent fixation on fibronectin-coated dishes (Labtek). Cells were fixed using 4% paraformaldehyde (PFA) (Sigma) and incubated at room temperature for 20 minutes. Cells were permeabilised using 0.2% TritonX in PBS for 5 minutes. 5% bovine serum albumin (BSA)/PBS was used to block non-specific binding for 30 minutes at room temperature. Cells were incubated with primary antibodies (*α*-tubulin-FITC Sigma 1:400) or fluorescent conjugated small molecules (DAPI 1:1000 Invitrogen, Phalloidin-TRITC 1:2000 Sigma) in 1% BSA/PBS for 1 hour at room temperature. Fixed samples were imaged on a Leica TCS SPE 2 microscope using 63x objective (ACS APO 63x oil NA1.3 DIC=E, coverslip correction 0.17).

### Image analysis

Images were processed and analysed using Fiji/ImageJ version 1.0 (Schindelin *et al*., 2012). Confocal fluorescent images are displayed as single plane or maximum intensity projections, as indicated in the figure legends. For cell shape analyses, outlines were manually segmented from bright-field or confocal images using the polygon selections tool in Fiji. Spindle angle analyses were obtained as the angle between the x-axis and a line drawn between the two spindle poles segmented from maximum projections of confocal images.

Images of overlaying cell outlines were produced using CellProfiler version 3.1.8 software (Carpenter *et al*., 2006) and overlay pipelines.

For calculations of the Jaccard similarity and dissimilarity indices, the standard equations (Jaccard, 1912) were implemented using custom-made software in Python using the NumPy (Harris *et al*., 2020), scikit-image (Van Der Walt *et al*., 2014) and SciPy libraries (Virtanen *et al*., 2020). To obtain the Jaccard index with shape contribution only, the areas of the cell outlines were made equal by uniformly scaling one of the images. The contribution of cell displacement was then removed by translating one image so that its centroid matched the centroid of the other image.

### Statistical analysis

Graphs were produced in Graphpad Prism (version 9.1.2). Bar charts and scatter plots show mean with error bars showing standard deviation. Stacked bars of summary data indicate mean percentages totalling 100% with error bars showing standard deviation. Where indicated, data was pooled from independent experiments, where N = number of replicates. The number of cells (n) analysed in each condition is indicated in parentheses on plots. Statistical tests were carried out in Graphpad Prism. All data sets were tested for normality using the D’Agostino-Pearson test for normality. For normal data sets, student T-tests were used to test whether differences were statistically significant. All non-normal data sets were analysed using Mann-Whitney two-tailed test. *p<0.01 **p<0.001 ***p<0.0001 ****p<0.00001.

### Data and code availability

All data and code is available upon request.

## Acknowledgements

We thank Guillaume Charras for help with the construction of tubulin-GFP labelled cell lines and Katarzyna Plak for help with CellProfiler overlay pipelines. We also thank Andrew Vaughn, Ki Ng and John Gallagher for microscopy support and Julian Downward for reagents. This project was funded by Cancer Research UK: SG was supported by a CRUK Clinical Student studentship (5369923), HKM and BB were supported by a CRUK Programme Grant (C1529/A17343) and by an EPSRC/CRUK multi-disciplinary award (C1529/A23335). HKM is currently supported by a Sir Henry Dale Fellowship jointly funded by the Wellcome Trust and the Royal Society (Grant Number 222575/Z/21/Z). SG, BB and HKM would also like to thank the MRC Laboratory for Molecular Cell Biology (MC_CF12266) where this work was done, and BB would like to thank the MRC Laboratory for Molecular Biology for support.

## Author contributions

SG, HKM and BB conceived and guided the project, and wrote the manuscript. TM and BB secured the funding and provided mentorship. SG and HKM performed the experiments. SG, TW and HKM performed the analysis. All authors reviewed and edited the manuscript.

## Supplementary Information

**S1.**
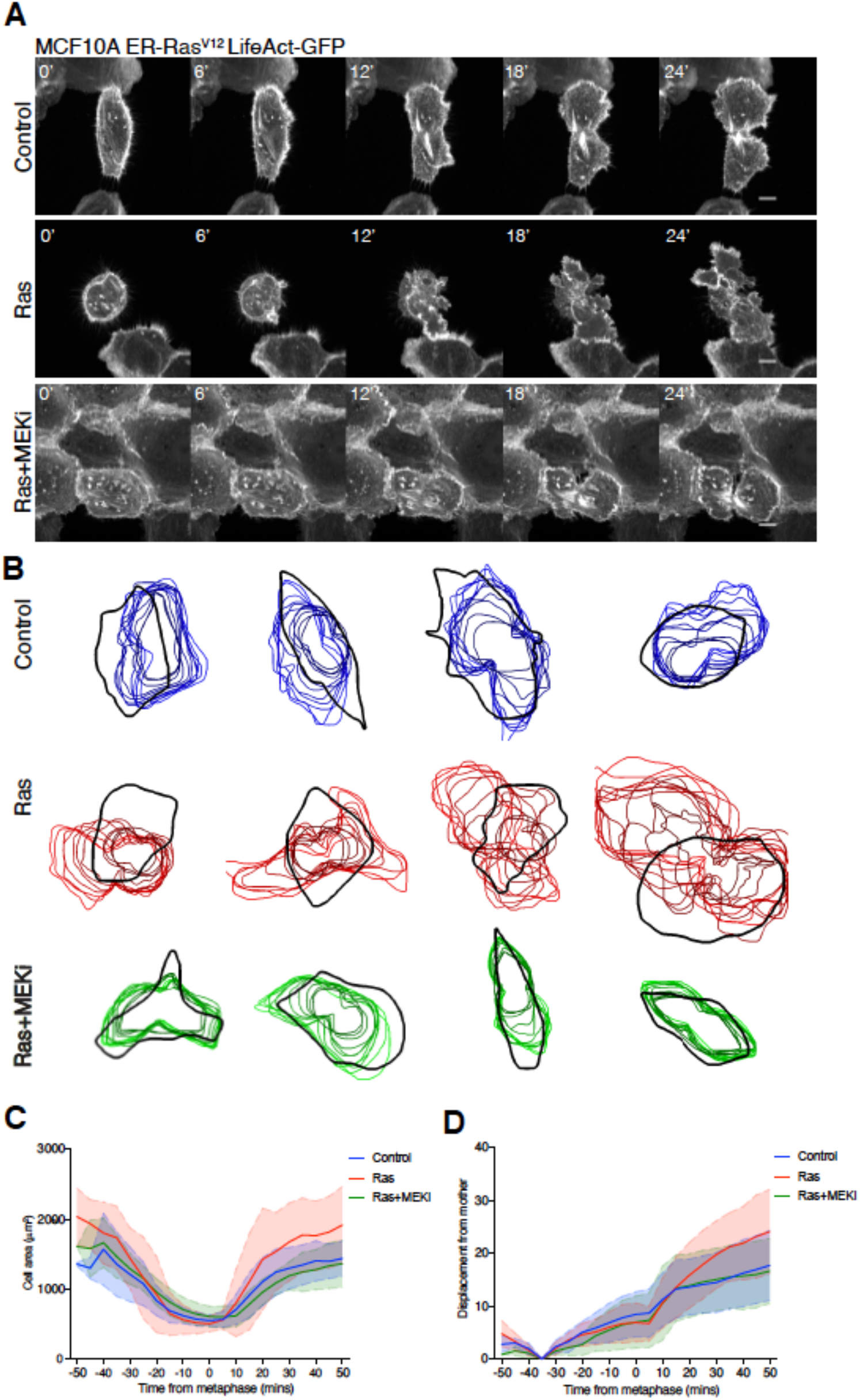
Analysis of post-mitotic respreading in MCF10A-LifeActGFP-ER-Ras^V12^ cells. (A) Time-lapse images of representative ER-Ras^V12^ cells labelled with LifeAct-GFP exiting mitosis. Cells were treated with a 5-hour treatment of ethanol, 4-OHT or 4-OHT + 10 μM selumetinib before imaging every 3 minutes. Images are presented as maximum projections. Time is aligned so t=0 represents the frame 3 minutes before anaphase. Scale bars represent 10μm. (B) Cell outlines as cells respread post-mitosis with respect to the mother cell shape. MCF10A-LifeAct-GFP-ER-Ras^V12^ cells were treated with ethanol, 4-OHT or 4-OHT plus 10μM selumetinib for 5-15 hours prior to time-lapse bright-field and fluorescence imaging every 5 minutes. Representative dividing cells were manually segmented at interphase (black), defined using the bright-field channel as 15 minutes before nuclear envelope breakdown (NEB). Post-mitotic respreading cell outlines were segmented at 5-minute intervals for 10 consecutive frames from the final frame of metaphase following ethanol (blue), 4-OHT (red) or 4-OHT plus MEKi (green). Image overlays were produced using CellProfiler software as described in Methods. (C) Graph to show the cell area of ER-Ras^V12^ cells treated with the conditions described in (B) as they progress through mitosis at consecutive time-points. n= 10 cells. (D) Graph to show centroid displacement of cells with respect to the mother of cells treated with the conditions described in (B) as they progress through mitosis. n= 10 cells.

**S2.**
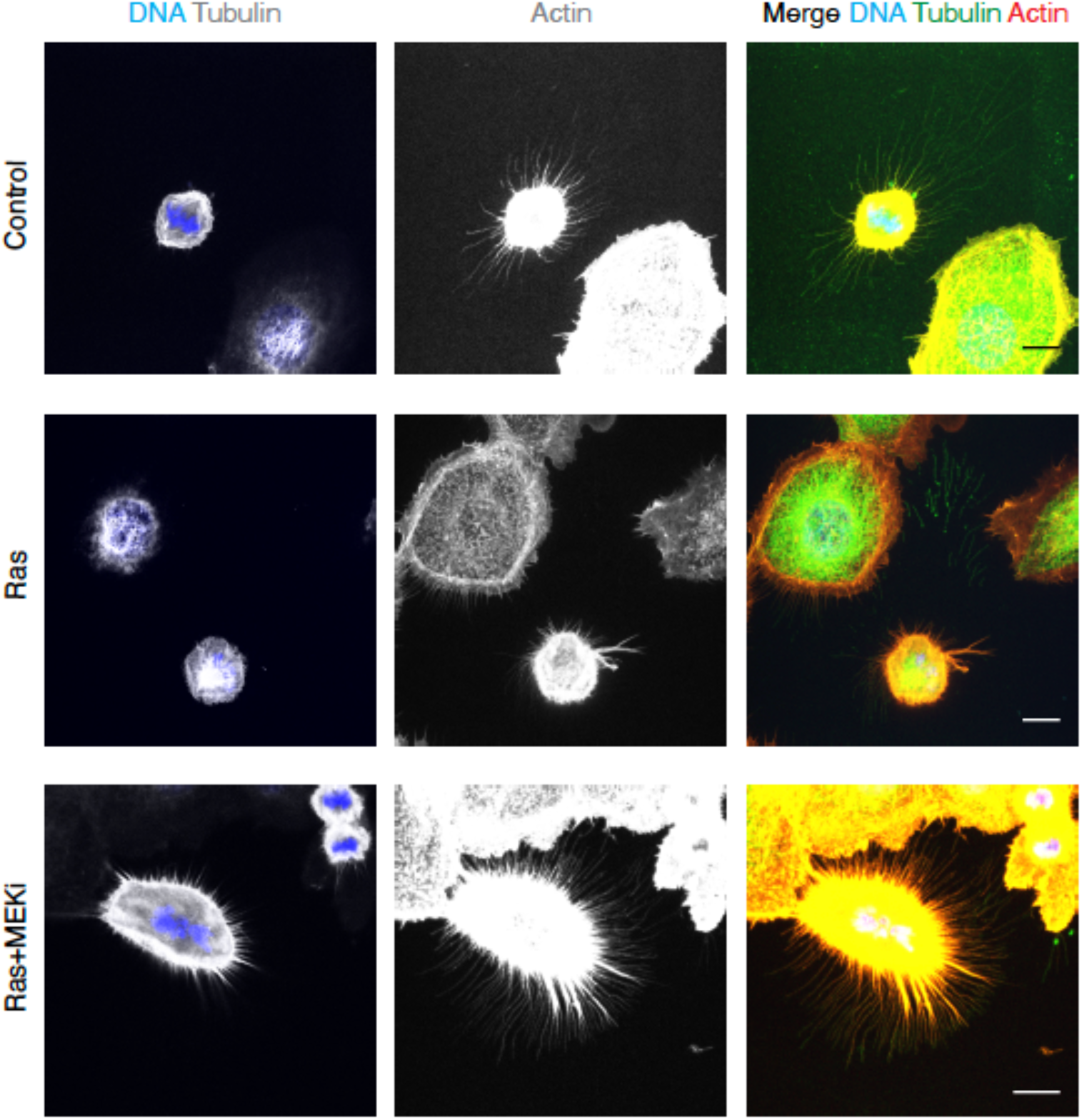
Immunofluorescence imaging of retraction fibres in MCF10A-ER-Ras^V12^ cells. Immunofluorescence images of MCF10A-ER-Ras^V12^ cells in metaphase following a 5-hour treatment of ethanol, 4-OHT or 4-OHT+ 10 μM selumetinib (MEK inhibitor). Cells are stained with DAPI, monomeric α-tubulin and phalloidin-TRITC. DAPI and tubulin images are displayed as a single z-slice, actin and merge images are displayed as maxiumum projections. Scale bars represent 10μm.

**S3.**
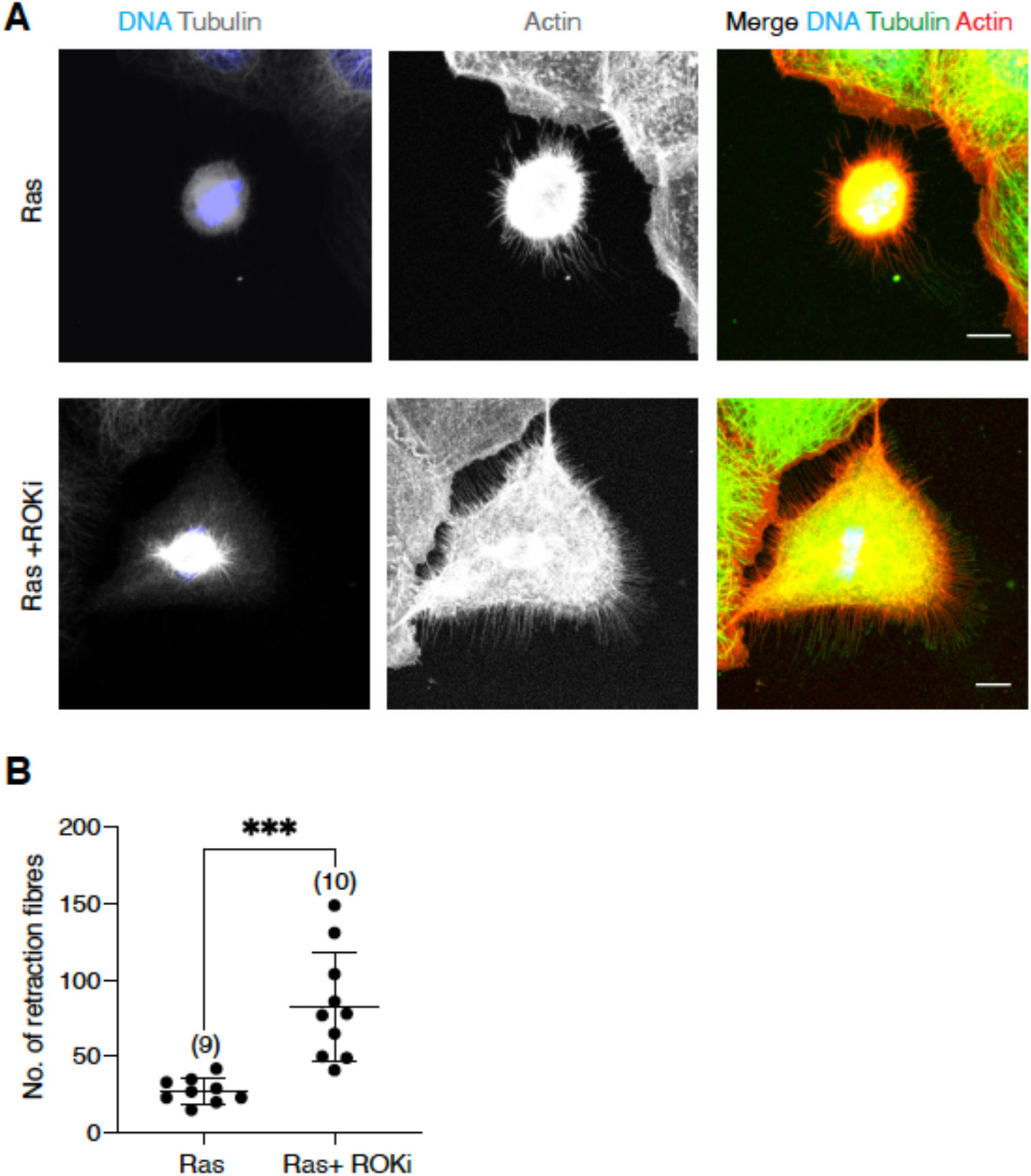
Immunofluorescence imaging of retraction fibres following ROCK inhibition. (A) Immunofluorescence images of MCF10A-ER-Ras^V12^ cells in metaphase following a 5-hour treatment of 4-OHT or 4-OHT+ 25μM Y27632 (ROCK inhibitor). Cells are stained with DAPI, monomeric α-tubulin and phalloidin-TRITC. DAPI and tubulin images are displayed as a single z-slice, actin and merge images are displayed as maxiumum projections. Scale bars represent 10μm. (B) Quantification of the total number of retraction fibres in single cells from the conditions described in (A).

**S4.**
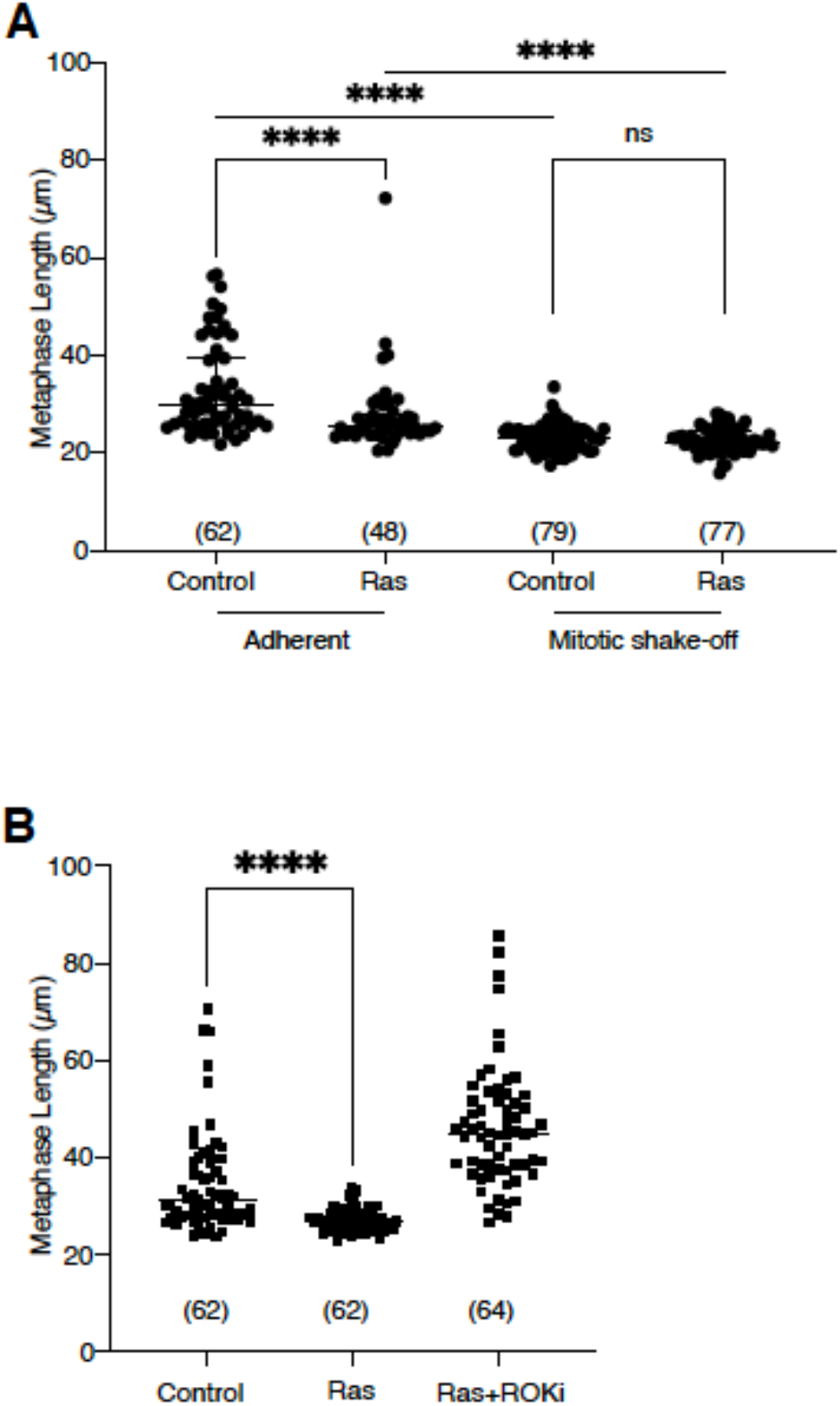
Metaphase cell length analysis during mitotic shake-off and ROCK inhibition. (A) Plot of individual metaphase cell length (Feret) measurements (taken as the frame 5 minutes before anaphase elongation) from the mitotic shake off experiment described in Fig. 4e. N= 3 experiments. (B) Plot of individual metaphase cell length (Feret) measurements from the ROCK inhibitor experiment described in Fig. 5c. N= 3 experiments.

